# The meiotic recombination landscape of *Drosophila virilis* is robust to mitotic damage during hybrid dysgenesis

**DOI:** 10.1101/342824

**Authors:** Lucas W. Hemmer, Guilherme Dias, Brittny Smith, Kelley Van Vaerenberghe, Ashley Howard, Casey M. Bergman, Justin P. Blumenstiel

## Abstract

Germline DNA damage is a double-edged sword. Programmed double-strand breaks establish the foundation for meiotic recombination and chromosome segregation. However, double-strand breaks also pose a significant challenge for genome stability. Because of this, meiotic double-strand break formation is tightly regulated. However, natural selection can favor selfish behavior in the germline and transposable elements can cause double-strand breaks independent of the carefully regulated meiotic process. To understand how the regulatory mechanisms of meiotic recombination accommodate unregulated transposition, we have characterized the female recombination landscape in a syndrome of hybrid dysgenesis in *Drosophila virilis*. In this system, a cross between two strains of *D. virilis* with divergent transposable element and piRNA profiles results in germline transposition of diverse transposable elements, reduced fertility, and male recombination. We sought to determine how increased transposition during hybrid dysgenesis might perturb the meiotic recombination landscape. Our results show that the overall frequency and distribution of meiotic recombination is extremely robust to germline transposable element activation. However, we also find that hybrid dysgenesis can result in mitotic recombination within the female germline. Overall, these results show that landscape of meiotic recombination may be insensitive to the DNA damage caused by transposition during early development.

## INTRODUCTION

Recombination during meiosis plays a critical role in ensuring proper chromosome segregation (Koehler *et al*. 1996). In addition, meiotic recombination benefits populations since it creates new genotypic combinations that can facilitate adaptation to changing environments (Weismann *et al*. 1904; Burt 2000). While meiosis and sexual reproduction is a conserved process across many eukaryotes, it can be exploited by selfish genetic elements. Transposable elements (TEs) are an example of one such class of selfish genetic element. TEs can increase within the genome, even to the detriment of the host, because fertilization provides a continuous opportunity for TEs to colonize new genomes (Hickey 1982) and recombination uncouples progenitor TEs from the harmful effects of descendant insertions. Upon proliferation, TEs can cause mutation, mis-regulation of gene expression, chromosomal rearrangement, and illegitimate recombination (Kidwell *et al*. 1977; Kazazian *et al*. 1988; Bennetzen 2000; Slotkin *et al*. 2007; Zhang *et al*. 2011). Moreover, TEs can activate the DNA damage response within developing germline stem cells and alter stem cell fate (Chen *et al*. 2007; Wylie *et al*. 2014; Tasnim and Kelleher 2018).

The harmful effects of TEs are especially evident in syndromes of hybrid dysgenesis, where sterility can arise in intraspecific crosses between males carrying TEs and females that lack them (Bingham *et al*. 1982; Bucheton *et al*. 1984; Yannopoulos *et al*. 1987; Lozovskaya *et al*. 1990). Hybrid dysgenesis is the result of TE activation in the absence of maternal repression by PIWI-interacting RNAs (piRNAs) (Aravin *et al*. 2007; Brennecke *et al*. 2008). The piRNA system of genome defense requires maternal deposition of piRNA to successfully silence TEs across generations. The combination of unrecognized TEs introduced to a naive genome *via* sperm and the absence of corresponding piRNAs in the egg results in TE activation and hybrid dysgenesis (Brennecke *et al*. 2008). A unique syndrome of hybrid dysgenesis in *D. virilis* is observed in intraspecific crosses between males of an inducing strain (designated strain 160) and reactive strain females (designated strain 9) (Lozovskaya *et al*. 1990). The primary TE family responsible for inducing dysgenesis remains unknown and sterility appears to be due to the mass activation of several TE families abundant in strain 160 but not strain 9. At least four elements are proposed to contribute significantly to dysgenesis: *Penelope*, *Helena*, *Paris*, and *Polyphemus* (Petrov *et al*. 1995; Evgen’ev *et al*. 1997; Vieira *et al*. 1998; Blumenstiel 2014; Funikov *et al*. 2018). In addition, activation of TEs during hybrid dysgenesis also leads to transposition of diverse TEs that are equally abundant between the two strains (Petrov *et al*. 1995).

The impact of hybrid dysgenesis on recombination during meiosis is an open question. TEs may modulate recombination rates directly since transposition itself can induce illegitimate recombination at the sites of DNA breaks. As a consequence, in the *P*-*M* system of hybrid dysgenesis in *D. melanogaster*, males undergo recombination (Hiraizumi 1971; Kidwell and Kidwell 1975; Kidwell *et al*. 1977). This is abnormal since meiotic recombination is absent in *D. melanogaster* males. However, rates of meiotic recombination induced by *P-M* dysgenesis are typically low, at a rate approximately 2-3% of female meiotic recombination (Preston and Engels 1996). *P-M* induced male recombination is usually attributed to an increased rate of mitotic exchange induced by DNA damage (Isackson *et al*. 1981). Many of these recombination events occur near *P* element insertion, require transposase, and are likely the byproduct of *P* element excision events that are repaired from the homolog (McCarron *et al*. 1994; Sved *et al*. 1995; Gray *et al*. 1996; Preston and Engels 1996; Preston *et al*. 1996). The effects of hybrid dysgenesis on female meiotic recombination rates in *D. melanogaster* are less clear. Changes in female recombination rates were not initially observed during *P-M* hybrid dysgenesis (Hiraizumi 1971; Chaboissier *et al*. 1995) but later studies found a slight increase (Kidwell 1977; Broadhead *et al*. 1977; Sved *et al*. 1991), often within pericentric heterochromatin. Slightly increased rates of female recombination were also observed within the *I*-*R* element system, but these were not localized near the centromere (Chaboissier *et al*. 1995). However, others have identified no increase in female recombination rates caused by *P-M* hybrid dysgenesis but, rather, a redistribution towards regions with low recombination and heterochromatic regions near the centromere (Slatko 1978; Hiraizumi 1981). There have been no investigations of how hybrid dysgenesis influences female meiotic recombination in *D. virilis*.

To determine how transposition influences recombination in *Drosophila* females, fine-scale genetic maps are required. *D. virilis* genetic maps have been obtained only with a limited number of markers which show that the rate of recombination in *D. virilis* is significantly higher than in *D. melanogaster* even though previously estimated rates also differ between studies (Weinstein 1920; Gubenko and Evgen’ev 1984; Huttunen *et al*. 2004) (Table 1). Here, we provide the first fine-scale genetic map for *D. virilis* using thousands of genotypic markers. Using this map, we investigate differences in both crossover (CO) frequency and distribution in the hybrid dysgenesis syndrome of *D. virilis*. Interestingly, we find no significant differences in meiotic recombination between genetically identical dysgenic and non-dysgenic female progeny. Instead, we find a strong signature of mitotic recombination that is likely derived from transposition events in the early germline. Thus, at least in *D. virilis*, meiotic recombination is remarkably robust to TE activation during hybrid dysgenesis. This suggests that the effects of DNA damage during early development are not sufficient to trigger changes in the control of recombination later during meiosis.

**Table 1:** Genetic map lengths of *D. virilis* chromosomes in centiMorgans reported in previous studies and this study.

|  | Chromosome |  |  |  |  |  |
| --- | --- | --- | --- | --- | --- | --- |
|  | X | 2 | 3 | 4 | 5 | 6 |
| Gubenko & Evgen'ev (1984) | 170 | 257 | 145 | 108 | 203 | 1 |
| Huttunen *et al.* (2004) | – | 118 | 125 | 147 | 60 | – |
| Current study | 143.5 | 160.9 | 139.6 | 148.3 | 140.0 | – |

## MATERIALS AND METHODS

### Fly Stocks and crosses

The hybrid dysgenesis syndrome in *D. virilis* is observed in crosses between reactive strain 9 females and inducer strain 160 males (Lozovskaya *et al*. 1990; Erwin *et al*. 2015). The study was performed in two stages. A smaller pilot study was performed, followed by a larger second study that incorporated additional optimization. We observed no differences between these two experiments, so we combined results for final analysis (see below). For both experiments, each strain and all subsequent crosses were maintained on standard media at 25°C. Prior to creating dysgenic and non-dysgenic hybrids, strain 160 and strain 9 were each inbred for 10 generations by single sib-pair matings. For each direction of the cross, approximately 20 4- to 6-day old virgin females of one strain and 20 2- to 10-day old males of the other strain were crossed *en masse* in bottles for six days. Strain 160 males crossed with strain 9 females induce dysgenesis in the F1 generation while the reciprocal cross yields non-dysgenic F1 progeny. Reciprocal crosses yield F1 files with identical genetic backgrounds, with the exception of the mitochondrial genome. By comparing the recombination landscape between F1 females with identical nuclear genotypes, we obtain a robust analysis of how hybrid dysgenesis influences recombination that effectively controls for genetic background. Individual virgin F1 females, four days post-emergence, were backcrossed in single vials to two or three 2- to 10-day old strain 9 males and maintained in vials for six days. Some non-dysgenic females were only allowed to lay for four or five days due to their high fertility to prevent vial crowding. Because fertility was low in dysgenic females, and to increase sample size of progeny within cohorts in the second experiment, a second brood was obtained from some dysgenic F1 females by transferring to another vial after ten days. These females were allowed to lay for an additional four days. We found no difference in recombination between first and second broods (see below), so results were combined across broods. Female backcross progeny (BC1) were collected once per day and immediately frozen at -20°C. Between 12 and 20 sisters from each non-dysgenic F1 female was collected as a sibship. All female progeny of the dysgenic F1 backcrosses were collected.

There is a high amount of variation in fecundity in dysgenic females. Some females are completely sterile, others have only reduced fecundity and some even have high numbers of progeny. One approach to analyzing the effects of dysgenesis on recombination would be to sample only single daughters from each F1 female. However, this approach would not allow for the discovery of recombination events arising as clusters within the germline. Therefore, we elected to sequence cohorts of BC1 sisters, balancing our sequencing across cohorts with different levels of fecundity. To determine if there might be an effect of fecundity on recombination, all male and female BC1 progeny across the two broods from the second larger experiment were counted to estimate the fecundity of the F1 mother. In some cases, dysgenic F1 females escape the effects of dysgenesis completely and produce as many progeny as non-dysgenic females. For these dysgenic F1 females, designated high fecundity, approximately 40 BC1 female progeny were subjected to recombination analysis by sequencing. Progeny produced by the low fecundity F1 dysgenic females were collected with cohort sizes ranging from one to 40 sisters. By sampling larger cohorts from high fecundity F1 dysgenic females, we sought to identify clustered recombination events derived within the germline of single females. Power to detect these events among a small cohort of sisters is low. By examining recombination in both high fecundity and low fecundity dysgenic females, we also obtained an additional comparison in the analysis of recombination landscapes between two outcomes of TE activation: TE activation with strong deleterious effects on fertility versus TE activation with no observable negative effects on fertility. For a full description of sampling, see Table S1.

### DNA extraction, library preparation, and Illumina sequencing

Sequencing libraries were prepared in two batches using different protocols. The first batch included 192 samples and library preparations were performed following the protocol outlined in Andolfatto *et al*. (2011) with minor modifications. Single flies were placed into a U-bottom polypropylene 96-well plate with lysis buffer and 3.5 mm steel grinding balls (BioSpec) then homogenized with a MiniBeadBeater-96 at 2,100 rpm for 45 seconds. DNA extractions on homogenized tissue were performed with the Agencourt DNAdvance Genomic DNA Isolation Kit (Beckman Coulter) following the Insect Tissue Protocol. DNA quantification was spot checked on some samples and estimated to average 1-2 ng/μl. For each sample, 10 μl of genomic DNA was digested with 3.3 U of MseI in 20 μls of reaction volume for four hours at 37°C, followed by heat inactivation at 65°C for 20 min. FC1 and FC2 adaptors (Andolfatto *et al*. 2011) (Table S2-S3) were ligated to the digested DNA with 1 U of T4 DNA ligase (New England Biolabs) in 50 μl of reaction volume at 16°C for 5 h and inactivated at 65°C for 10 minutes. The samples were pooled and concentrated using isopropanol precipitation (1/10 vol NaOAc at pH 5.2, 1 vol of 100% isopropanol, and 1 μl glycogen). The library was resuspended in 125 μl of 1X Tris-EDTA (pH 8). Adapter dimers were removed with 1.5X vol AMPure XP Beads (Agencourt) and ligated products were resuspended in 32 μl of 1X Tris-EDTA (pH 8). 200-400 bp DNA fragments were selected using a BluePippin (Sage Science). Size-selected fragments were cleaned using 2X vol of AMPure XP beads and resuspended 20 μl of 1X elution buffer (10 μM Tris, pH 8). Libraries were quantified using a Qubit fluorometer before an 18-cycle PCR amplification on bar-coded fragments with Phusion high-fidelity PCR Kit (New England Biolabs). The PCR products were cleaned using 1X vol of AMPure XP Beads.

**Table 2:** Interference values (*nu*) and frequency of crossovers created in the non-interference pathway (*P*) for all chromosomes. Both values were estimated with the Housworth-Stahl model for the entire dataset.

|  | Chromosome |  |  |  |  |
| --- | --- | --- | --- | --- | --- |
|  | **X** | **2** | **3** | **4** | **5** |
| $Nu$ | 3.22 | 3.17 | 2.68 | 3.09 | 3.37 |
| $P$ | 4.81E-02 | 1.05E-02 | 2.73E-08 | 6.91E-03 | 9.75E-03 |

For the larger second batch (768 samples), sequencing libraries were constructed with an optimized rapid DNA extraction protocol that uses an in-house Tn5 transposase similar to the procedure outlined in Picelli *et al*. (2014). DNA was extracted using the *Quick*-DNA 96 kit (Zymo) and 1-2 ng of DNA was tagmented with Tn5 transposase stored at a concentration of 1.6 mg/ml with pre-annealed oligonucleotides. Tagmentation was performed in 20 μl reaction volumes containing 2 μl of 5X TAPS-DMF buffer (50 mM TAPS-NaOH, 25 mM MgCl2 (pH 8.5), 50% v/v DMF) and 2 μl of 5x TAPS-PEG buffer (50 mM TAPS-NaOH, 25 mM MgCl2 (pH 8.5), 60% v/v PEG 8000). Samples were incubated at 55°C for 7 min then rapidly lowered to a holding temperature of 10°C. Reactions were inactivated with 5 μl of 0.2% SDS followed by an additional incubation at 55°C for 7 min. PCR-based barcoding was performed on 2.5 ul of sample tagmentation volumes using the KAPA HiFi HotStart ReadyMix PCR Kit (Thermo Fisher Scientific), 1 μl of 4 μM Index 1 (i7) primers (Table S4), and 1 μl of 4 μM Index 2 (i5) primers (Table S5) in 9 μl of reaction volume. The PCR thermocycling conditions were: 3 min at 72°C, 2 min 45 sec at 98°C, followed by 14 cycles of 98°C for 15 sec, 62°C for 30 sec, 72°C for 1 min 30 sec. PCR-amplified samples were pooled and the pooled samples were cleaned using 0.8 X vol AMPure XP Beads. We size-selected DNA fragments 250-400 bp from the pooled sample on a BluePippin and cleaned using 1X vol of AMPure XP Beads.

**Table 3:** Pearson’s correlation coeffecients (*R*) and *p*-values between rates of recombination and sequence parameters calculated in 250 kb intervals.

| Sequence Parameter |  | Chromosome |  |  |  |  | Total | Total minus<br>Zero CO |
| --- | --- | --- | --- | --- | --- | --- | --- | --- |
|  |  | X | 2 | 3 | 4 | 5 |  |  |
| GC Content | R | 0.08 | 0.35 | 0.11 | 0.18 | 0.33 | 0.23 | 0.15 |
|  | *p* | 0.372 | <0.001 | 0.263 | 0.079 | <0.001 | <0.001 | <0.001 |
| Gene density | R | 0.31 | 0.21 | 0.12 | 0.32 | 0.33 | 0.19 | 0.03 |
|  | *p* | <0.001 | 0.012 | 0.222 | <0.001 | <0.001 | <0.001 | 0.506 |
| Simple repeats | R | 0.44 | 0.43 | 0.31 | 0.32 | 0.54 | 0.39 | 0.177 |
|  | *p* | <0.001 | <0.001 | <0.001 | <0.001 | <0.001 | <1E-10 | <0.001 |
| SNP Density | R | 0.64 | 0.553 | 0.60 | 0.67 | 0.65 | 0.62 | 0.49 |
|  | *p* | <1E-10 | <1E-10 | <1E-10 | <1E-10 | <1E-10 | <1E-10 | <1E-10 |
| TE Density | R | -0.47 | -0.47 | -0.33 | -0.44 | -0.49 | -0.44 | -0.14 |
|  | *p* | <0.001 | <0.001 | <0.001 | <0.001 | <0.001 | <1E-10 | <0.001 |

**Table 4:** Clusters of recombination identified in the BC1 progeny.

| F1 Female Identifier | Chromosome | Position (Mb from Telomere) | CO Progeny / Total Progeny | Distortion Genotype | TEs detected in Strain 160 |
| --- | --- | --- | --- | --- | --- |
| 3* | X | 17.7 | 4 / 29 | None | *Polyphemus* |
| 111 | 3 | 25.1 | 4 / 11 | Strain 9** | *Polyphemus* |
| 4013 | X | 11.4 | 4 / 13 | Strain 160 | None |
| 4029 | X | 21.3 | 14 / 19 | Strain 160 | None |
| 5011 | 3 | 9.8 | 28 / 32 | None | *Polyphemus* |
| 5019 | 3 | 20.2 | 10 / 30 | Strain 9** | None |
| 5022 | 3 | 23.6 | 5 / 11 | Strain 9** | *Paris* |
| 5089 | X | 23.4 | 4 / 11 | Strain 160 | *Polyphemus* x2, *Paris* |

\* The F1 female is non-dysgenic
\*\* Segregation distortion occurred in some of the progeny. The rest appear normal and probably
did not inherit a mitotic CO chromatid

All libraries were sequenced at the University of Kansas Genomics Core on an Illumina HiSeq 2500 Sequencer with 100 bp single-end sequencing. The first 192 samples were sequenced on two lanes using the Rapid-Run Mode while the Tn5-produced libraries were sequenced on two lanes using the High-Output Mode (summary of the output is in File S2).

### DNA extraction, library preparation, Pacbio sequencing and assembly

PacBio sequencing was performed on strain 160 after 10 generations of single-sib mating followed by re-validation for induction of dysgenesis. Females collected for DNA extraction were allowed to eclose over 10 days, aged for two additional days, starved for 12 hours to evacuate the gut, then immediately frozen in liquid nitrogen. 500 mg of whole flies were then used for DNA extraction with Blood Cell and Culture Midi Kit (Qiagen) (Chakraborty *et al*. 2016). The mortar was pre-chilled with liquid nitrogen prior to grinding and the resulting fine powder was directly transferred into Buffer G2. DNA extraction was performed across 5 columns, using a total of 47.5 mls G2, 190 μl RNAse A (100 mg/ml) and 1.25 ml of Protease from the Qiagen Kit. 50 mls were placed in a 50°C shaker overnight. After overnight incubation, debris was spun down and poured onto column. The elution was performed according to manufacturer’s instructions and precipitated with 0.7 volumes of isopropanol, followed by spooling with a glass rod and resuspending in 100ul EB Buffer. The final DNA concentration was estimated with a Qubit to be 843 ng/ul, yielding approximately 85 μg. PacBio sequencing was performed by the University of Michigan DNA Sequencing Core.

Purified strain 160 DNA was used to generate PacBio SMRTbell libraries using the protocol: Procedure & Checklist 20 kb Template Preparation using BluePippin Size Selection. Briefly, approximately 10 μg of template was sheared using Covaris g-TUBES to obtain a 20-25 Kb target length. Sheared DNA was purified using pre-washed AMPureXP beads, analyzed for size and concentration, then treated with Exo VII to remove single stranded DNA, followed by DNA damage and end repair. End repaired DNA was then blunt ligated to adaptors to form SMRTbells and treated with Exo VII plus Exo III to remove any fragments that lack adaptors on both ends.

SMRTbells were purified using pre-washed AMPureXP beads, analyzed for size and concentration, then run through a Sage Scientific Blue Pippin instrument with 0.75% agarose dye-free cassette and S1 external marker to remove templates smaller than 10 kb. The PacBio Binding Calculator was used to determine conditions to anneal primers to SMRTbells and bind DNA polymerase (P6/C4 chemistry) to form SMRTbell-primer-polymerase complexes.

Complexes were bound to Magbeads and washed to remove unbound template and other contaminants. Purified complexes with an added Pacific Biosciences internal control were loaded on a PacBio RS II instrument and run using the MagBead-OCPW protocol. The resulting library was sequenced on 21 SMRT cells with a movie time of 360 minutes per SMRT cell, totalling ∼80-fold coverage of the estimated ∼380 Mb *D. virilis* genome (Bosco *et al*. 2007).

Assembly of the PacBio reads was performed using Canu v1.5 with default settings (Koren *et al*. 2017). Canu performs read correction on the 40x longest reads based on the genomeSize parameter. This parameter was set to 200 Mb after analyzing the read size distribution to avoid including shorter reads that could result in deterioration of assembly quality. The raw PacBio reads were aligned back to the Canu assembly with pbmm2 v1.0.0. and the assembly was polished with Arrow using gcpp v0.01.e12a6d6. PacBio polishing software were downloaded as part of the pb-assembly metapackage v0.0.6 from bioconda (https://github.com/PacificBiosciences/pb-assembly). A second round of polishing was performed after aligning Illumina reads from strain 160 (SRR1200631, Erwin *et al*. 2015) with BWA-MEM v0.7.17-r1188 (Li 2013) and correcting errors with Pilon v1.22 (Walker *et al*. 2014). Since *D. virilis* strain 160 is largely colinear with the current *D. virilis* reference genome (strain 15010-1051.87; *Drosophila* 12 Genomes Consortium 2007), we performed reference-based scaffolding of the strain 160 PacBio assembly using RaGOO v1.1 (Alonge *et al*. 2019). RaGOO performs scaffolding and correction of potential misassemblies by aligning *de novo* assemblies to a reference genome and breaking chimeric contigs where necessary. For the reference, we used an updated version of the SNP-corrected reference genome for strain 160 generated in Erwin *et al*. (2015) that incorporates the chromosome mapping data generated by Schaeffer *et al*. (2008; see below). General assembly statistics were analyzed with QUAST v5.0.2 (https://github.com/ablab/quast, commit 67a1136, Gurevich *et al*. 2013). Assembly completeness was assessed by searching for highly conserved orthologs with BUSCO v3.0.2 (Simão *et al*. 2015) using the Diptera ortholog gene set from OrthoDB v9 (Zdobnov *et al*. 2016). Assembly statistics are available in Table S6. Coordinates of the mitotic CO clusters (see methods below) were lifted over to the final version of the PacBio assembly using minimap2 2.16-r922 (Li 2018).

### Annotation of genome resources

Illumina-based reference genomes for strains 9 and 160 (Erwin *et al*. 2015) based on the original Sanger shotgun sequence assembly of *D. virilis* (*Drosophila* 12 Genomes Consortium 2007) were annotated with the most up-to-date gff file for *D. virilis* (v1.6 Flybase, Gramates *et al*. 2017) using Maker v3.31.9 on default settings (Cantarel *et al*. 2008). Due to errors in the original reference assembly, genome region 33,464,439-35,498,860 on Chromosome 2 was excluded and genome region 22,673,797-24,330,628 on Chromosome 5 was placed at position 3,761,384 on the same chromosome. Thus, previous strain-specific reference genomes (Erwin *et al*. 2015) were adjusted for two mis-assemblies and updated as ‘_2’ (strain 9, File S3; strain 160, File S4). TE annotations for Illumina-based reference genomes were obtained using RepeatMasker v4.06 (Tarailo-Graovac and Chen 2009) with the custom TE library from Erwin *et al*. (2015). TE annotation of the strain 160 PacBio assembly was also obtained using RepeatMasker with the custom TE library from Erwin *et al*. (2015).

### Crossover quantification

Illumina FASTQ files were parsed according to barcode sequences and trimmed by the University of Kansas Genomics Core facility. The FASTQ files were mapped to the Illumina-based reference genomes for strains 9 and 160 using the multiplex shotgun genotyping (MSG: https://github.com/JaneliaSciComp/msg) bioinformatic pipeline for identifying reliable markers and determining ancestry at those markers using a hidden Markov model (HMM) v0.1 (Andolfatto *et al*. 2011). Briefly, reads were mapped with BWA aln to the strain 9 and 160 parental reference _2 files. Output files were used for HMM determination of ancestry along the length of the chromosomal segments (see control file, File S5, for settings). The MSG pipeline provides both ancestry probability calls and CO positions, along with an estimate of the boundaries for CO positions. Samples with fewer than 10,000 reads were discarded. Double COs less than 750 kb apart were discarded as these events were considered unlikely. We observed tha reads mapping to regions near the telomere and centromere often predicted the same genotype across all samples. In principle, this could be driven by segregation distortion. However, these regions also showed low density for uniquely mapped reads. In addition, segregation distortion for these regions would drive distortion of linked flanking markers, which we did not see.

Therefore, we considered these regions problematic and removed them from the analysis. COs located within 500 kb of the telomere of the X and 4th chromosome and COs within 700 kb on the 2nd, 3rd, and 5th chromosomes were removed. COs near the centromeres were also removed as follows: within 3.5 Mb on the X chromosome, within 1.1 Mb on the 2nd chromosome, within 1.5 Mb on the 3rd chromosome, within 2.4 Mb on the 4th chromosome, and 2.3 Mb on the 5th chromosome. The 6th chromosome (corresponding to the 4th in *D. melanogaster*) was also removed from analysis. In addition, we performed additional manual curation of COs to remove calls that appeared incorrect. For example, double COs that were spaced closely in samples with lower numbers of reads and based on ancestry probabilities that were less than 0.9

## Data analysis

CO outputs from the MSG pipeline were analyzed with R Version 3.4.2 (R Core Team 2017). Ancestry probability outputs were used in the R/qtl package (Broman *et al*. 2003) to construct genetic maps and calculate interference using the Housworth Stahl model (Housworth and Stahl 2003) using the xoi package (Broman and Kwak, Version 0.67-4). We used the lme4 (Bates *et al*. 2015) and lsmeans (Lenth 2016) packages for mixed-model testing of CO events in BC1 progeny. The model construction was performed using the glmer() and glm() functions to test the random effects of F1 female and fecundity of the F1 female and the fixed effects of dysgenesis and batches. Fecundity estimates obtained from dysgenic crosses in the second experiment were first used to determine if fecundity had an effect on total CO count. We found that fecundity had no effect on CO count (χ^2^1= 2.02, *p* = 0.155) and excluded it from the final model.

The model for how these effects predict total CO numbers in R was as follows:

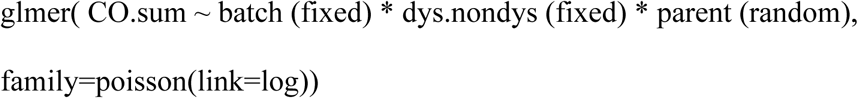

We used likelihood ratio tests to determine the significance of each effect on variance in total CO number. We used the Biostrings R package (Pagès et al. 2017) to analyze genomic sequences for correlations between genomic features and recombination. Figures were constructed using ggplot2 (Wickham 2016).

### Analysis of mitotic recombination

Mitotic (or pre-meiotic) recombination is identified by the presence of crossovers that are common among the progeny of a single parent. These are commonly designated as recombination clusters and are distinct from hotspots because they are found only within cohorts of siblings. Clusters indicating germline mitotic recombination were identified as COs in four or more progeny of a single F1 mother within a 100 kb span. Since the fecundity effects of hybrid dysgenesis are highly variable, there was imbalance in progeny counts from dysgenic and non-dysgenic backcrosses. It was therefore necessary to account for this variation in the estimation of rates of mitotic recombination. This was achieved using a likelihood approach to determine if rates of mitotic cluster formation (α) within the germline and the frequency of mitotic recombination within cohorts (β, conditional upon cluster formation) differed between dysgenic and non-dysgenic parents. Only one mitotic cluster was ever observed per single chromosome so rate estimate was performed on a per chromosome basis. The probability of not observing a cluster event (on a given chromosome) is given by the probability that a mitotic recombination event does not occur in the germline (1-α) plus the probability that a mitotic recombination event does occur (α) but is not observed among the sampled progeny:

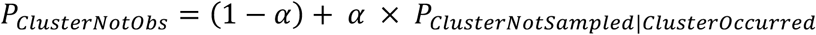

Conditional on mitotic recombination occurring, the probability that it was not observed is equal to the probability that three or fewer progeny within the cohort inherit the recombinant chromatid from the mother. This is given with the binomial probability distribution where β is the frequency of recombinant chromosomes transmitted by the mother with the mitotic event:

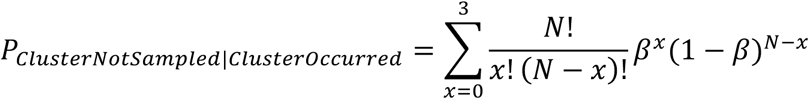

where *N* is the total number of progeny in the cohort, β and is the frequency of progeny that inherit the recombinant chromosome. Therefore, parents with three or fewer progeny have

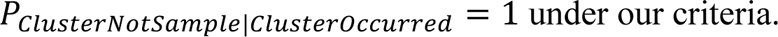

When a cluster event is observed, the probability of *x* progeny with the recombinant chromosome is given by:

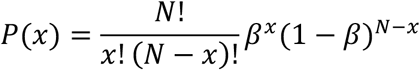

Overall, the probability that a cluster is observed at a given frequency within a cohort is equal to the probability that mitotic recombination happened (α) multiplied by the probability that it is observed at a given frequency, conditional on it having happened:

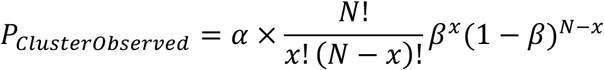

The full likelihood of the data is thus given by:

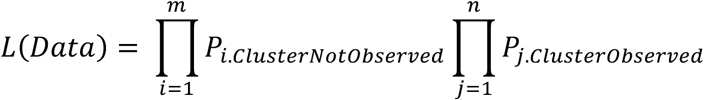

where *i* is index of mothers without an observed mitotic cluster and *j* as the index of mothers whose progeny indicate a mitotic cluster. Taking the logarithm of our likelihood equation gives

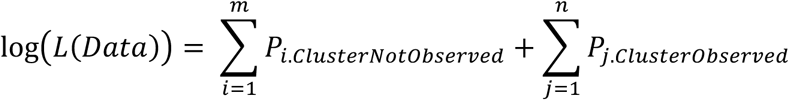

Mitotic recombination was only ever observed on the X and 3rd chromosomes so a combined rate was only estimated for these two chromosomes. To estimate rates of mitotic recombination across dysgenic and non-dysgenic females, we used the Python module Scipy to maximize the log-likelihood of the data based on α and β. Nested likelihood ratio tests were used to determine whether there was support for unique values of α or β in dysgenic or non-dysgenic females. Two three-parameter models were used with distinct cluster formation rates for dysgenic (D) and non-dysgenic (ND) females (α_D_, α_ND_, β) and, reciprocally, separate frequencies of transmission of the recombinant chromatid (α, β_D_, β_ND_). We also used as a four-parameter model with individual estimates for the dysgenic and non-dysgenic mothers (α_D_, α_ND_, β_D_, β_ND_). 95% confidence intervals (95% CIs) for parameter estimates were obtained by determining parameter values with likelihood scores two log-likelihood units from the ML estimate with other maximum likelihood estimated parameters fixed. We tested if models were significantly improved with the inclusion of additional parameters with a likelihood ratio test (LRT) and a chi-squared distribution with one degree of freedom for every additional parameter estimated. The Python script for the maximum likelihood analysis of the mitotic recombination rates is in File S6.

## Data availability

All of the de-multiplexed Illumina sequencing reads from BC1 progeny, PacBio reads for strain 160, and the strain 160 PacBio assembly generated in this study are available at the National Center for Biotechnology Information under accession PRJNA553533. Supplemental files are available at FigShare.

## RESULTS

### Crossover detection by sequencing

To identify recombination events in reciprocal F1 dysgenic and non-dysgenic progeny, F1 females were backcrossed to reactive strain 9. By sequencing backcross (BC1) progeny, we identified the recombination events that occurred under the dysgenic and non-dysgenic condition in the germline of F1 females. F1 dysgenic and non-dysgenic female progeny have identical nuclear genotypes, which enables a controlled comparison of the effects of TE mobilization on the recombination landscape. In total, 828 BC1 female progeny were sequenced at sufficient depth to map recombination breakpoints; 132 samples had fewer than 10,000 reads and were not included in our analysis. 275 BC1 progeny were sequenced from 20 F1 non-dysgenic females, 311 BC1 progeny were sequenced from 66 low fecundity F1 dysgenic females, and 242 BC1 progeny were sequenced from seven high fecundity F1 dysgenic females. Across all samples, the MSG pipeline identified a total of 1,150,592 quality-filtered SNPs between the two parental genomes with an average of distance of 136 bp between SNPs. The MSG HMM uses relative mapping abundance of reads that are uniquely derived from one of the two parental strains.

Using this information, combined with the crossing scheme, it provides genotype probabilities for each SNP. For each sample, and at each SNP, the HMM provided a genotype probability of the BC1 progeny sample being either homozygous for strain 9 (the strain that the F1 progeny were backcrossed to) or heterozygous. CO breakpoint intervals were then defined based on local genotype probability calls that switch from greater than 95% to less than 5% along the chromosome. The median CO breakpoint interval calculated by the MSG HMM was approximately 18 kb and 84% of all COs localized within a span of 50 kb or less. Seventeen CO breakpoint intervals were approximately ∼ 1 Mb but those were in samples with low read counts near the 10,000 read cutoff for samples allowed in the analysis.

### A high-resolution genetic map of *D. virilis*

Previous studies indicate that the genetic map of *D. virilis* is approximately three times larger than the genetic map of *D. melanogaster* (Gubenko and Evgen’ev 1984; Huttunen *et al*. 2004).

Critically, the map lengths obtained in those two studies are quite different (Table 1), perhaps due to a limited number of genetic markers available. Our combined sample has a sufficient density of markers to provide the first high-resolution recombination map for *D. virilis*. Combining results from all samples to construct our map was reasonable, since the effects of dysgenesis were non-significant (see below).

The total genetic map length of *D. virilis* estimated in our combined sample is 732 cM (centiMorgans) or 2.5 times longer than the genetic map length of *D. melanogaster* (Comeron *et al*. 2012) (Table 1). The genetic map length estimated in the current study is more than 100 cM shorter than the first detailed genetic map of *D. virilis* (Gubenko and Evgen’ev 1984) (Table 1). This may be partly explained by our stringent exclusion of problematic genomic regions.

However, comparing chromosomes that were characterized in all three studies (2, 3, 4, and 5), our estimate for cM is within or very close to the upper estimate of the two prior studies. In addition, our cM estimates were more uniform across the chromosomes, which are all fairly similar in physical length. As expected, the genetic map length of each chromosome in our study correlates with physical length (*R^2^* = 0.851, *p* = 0.025). There is no significant correlation in the two prior studies (*R^2^* = 0.079, *p* = 0.72, Huttunen *et al*. 2004; *R^2^* = 0.358, *p* = 0.28, Gubenko and Evgen’ev 1984, excluding the 6^th^ chromosome). The differences in recombination rates between *D. virilis* and *D. melanogaster* might appear explained due to their differences in genome size. The estimated genome size of *D. virilis* is roughly twice the size of the *D. melanogaster* genome, (404 vs 201 Mb, Bosco *et al*. 2007). Thus, across the entire genome, the average rate of recombination in *D. virilis* is 1.8 cM/Mb and similar to the average recombination rate of 1.4 cM/Mb in *D. melanogaster*. However, close to half of the *D. virilis* genome is comprised of satellite DNA, with large expanses of centromeric heterochromatin where little or no crossing over takes place (Gall and Atherton 1974; Bosco *et al*. 2007). Thus, the *D. virilis* assembly of euchromatin, where most COs take place, is 206 Mb in length. Accounting for just euchromatic regions in both species, the average rate of recombination in euchromatin in *D. virilis* is twice as high as *D. melanogaster* based on euchromatic assembly genome size (3.5 cM/Mb vs. 1.8 cM/Mb). One possible reason for a higher rate of recombination in *D. virilis* may be the fact that pericentric heterochromatin comprised of satellite DNA may shield the chromosomes arms from the suppressive centromere effect (Hartmann *et al*. 2019).

Interference reduces the probability of an additional CO in proximity to other COs. We calculated interference in *D. virilis* using the Housworth-Stahl model to calculate *nu*, a unitless measure of interference, with a maximum likelihood function based on intercrossover distances (Housworth and Stahl 2003). If COs are not subject to interference, intercrossover distances are Poisson distributed and *nu* is equal to one (Broman and Weber 2000). *nu* values greater than one are observed when COs are spaced more evenly than expected under a Poisson process. Each chromosome in *D. virilis* has detectable interference between COs with an average *nu* ∼ 3 (Table 2). The Houseworth-Stahl model also estimates the percentage of COs produced through an alternative pathway not subject to interference as the escape parameter *P* (de los Santos *et al*. 2003). Less than 1% of the COs in our study are estimated to be produced through the alternative CO pathway (Table 2). In contrast to interference, assurance is the recombination control mechanism that maintains a minimal number of COs on each chromosome to ensure proper chromosome segregation during meiosis. In the absence of CO assurance and interference, the distribution of COs should resemble a Poisson distribution with variance in CO number equal to the mean. We find the CO mean and variance are not equal (7.3 and 4.9, respectively) and the distribution of CO counts is significantly different from a Poisson distribution (χ^2^ = 53.6, *p* = 5.74E-08). Moreover, *D. virilis* has a higher than expected frequency of individuals with CO numbers close to the mean (5-8 COs) and a lower than expected frequency of extreme CO numbers (Figure S1). This indicates the collective action of CO assurance and interference.

In many organisms, the total number of COs between markers in a single tetrad is unobservable because COs are typically detected using random spore analysis based on only one of the four chromatids resulting from meiosis. In contrast, tetrad analysis uses the number of COs for each of the four meiotic products to estimate the frequency of non-exchange tetrads (*E*_0_), single-exchange tetrads (*E*_1_), or multiple-exchange tetrads (*E*n) (Weinstein 1918). We used the Weinstein method to estimate the frequency of *E*_0_ tetrads for each chromosome in *D*. *virilis*. The X chromosome and third chromosome *E*_0_ tetrad frequencies were estimated at 1.2% and 2.1% respectively. These calculated *E*_0_ tetrad frequencies in *D. virilis* are lower in comparison to *D. melanogaster*, previously estimated to be 5-10% (Zwick *et al*. 1999; Hughes *et al*. 2018).

However, the Weinstein method indicated negative *E*_0_ tetrad frequencies ranging from -2.6% to −3.9% for the second, fourth, and fifth chromosomes. Negative values were also sometimes obtained for other exchange class frequencies (Table S7). Negative tetrad frequencies are a drawback to using the classic Weinstein method (Zwick *et al*. 1999). Nonetheless, these results indicate that there are many fewer non-exchange tetrads in *D. virilis* compared to *D. melanogaster*, consistent with the higher estimated rate of recombination per Mb in *D. virilis*.

Recombination rates are often correlated with certain sequence features, such as GC content, simple motifs, and nucleotide polymorphism (Begun and Aquadro 1992; Kong *et al*. 2002; Comeron *et al*. 2012). In *D. virilis*, recombination rates appear to be weakly correlated with GC content and gene density as not all chromosomes show significant correlations with either sequence parameter (Table 3). This may be due to unusual patterns of recombination along the length of the chromosome (discussed later). Simple repeats and SNP density between the two strains show significant positive correlations with recombination rate on all chromosomes even after removal of non-recombining regions. Nucleotide diversity is frequently correlated with recombination rates (Begun and Aquadro 1992; Kong *et al*. 2002) and the strong correlation between SNP density and recombination in our data is consistent with this pattern (Figure S2). TE density shows a strong negative correlation with recombination rate but this becomes weaker when non-recombining regions are removed (Table 3, Figure S2). The similar pattern of weak correlation between TE density and recombination when regions without recombination are removed is also seen in *D. melanogaster* (Rizzon *et al*. 2002), where TEs mostly accumulate in non-recombining pericentromeric heterochromatin (Bartolomé *et al*. 2002; Bergman *et al*. 2006).

### No modulation of recombination rates during hybrid dysgenesis

To compare recombination rates in dysgenic and non-dysgenic females, we constructed a full mixed-effects likelihood model using the lme4 R package (Bates et al. 2015). The direction of the cross (dysgenic vs. non-dysgenic) and F1 collection batch (pilot vs. second experiment) were treated as fixed effects; the F1 female of origin was treated as a random effect. In the full model, we find no difference in the total number of COs between the pilot and second experiment (χ^2^ 1 = 0.10, *p* = 0.755). This suggests that there were no effects of library construction procedure and justifies combining data sets. Likewise, no effect was found for fecundity in dysgenic flies on CO numbers (χ^2^ 1 = 2.02, *p* = 0.155). Importantly, dysgenesis did not have a significant effect on total CO numbers (χ^2^ 1 = 0.45, *p* = 0.506) with nearly identical means in CO number between dysgenic and non-dysgenic progeny (Figure 1A). There was a marginal interaction between dysgenesis and batch (χ^2^ 1 = 3.17, *p* = 0.075), but this appears driven by a single high fecundity dysgenic F1. This F1 female, designated 701, revealed a significantly larger mean CO number than the other high fecundity dysgenic females (8.3 COs, least squares mean contrast, *p* = 0.021, Figure 1B). Without the 701 female, the interaction between dysgenesis and batch is non-existent (χ^2^ 1 = 0.06, *p* = 0.803). Overall, full model revealed that the identity of the female had minimal effect on CO number (variance < 0.0001).

**Figure 1:**
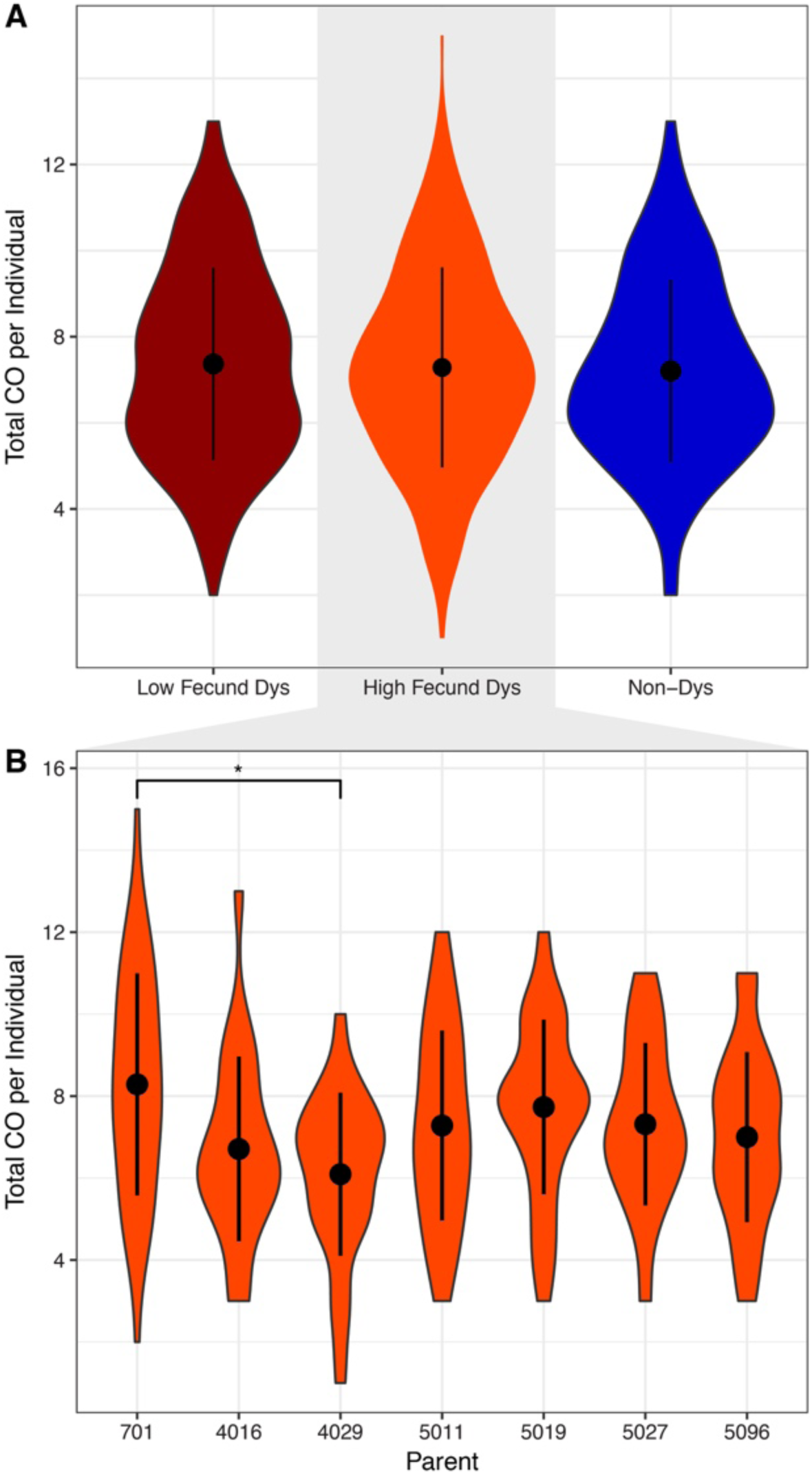
The distribution of the total crossover (CO) count observed in individual BC1 progeny with the mean and standard deviation. The mean for each group is represented with a black dot and the standard deviation is the black line. A) The distribution of the total CO count per BC1 progeny of low fecundity dysgenic, high fecundity and non-dysgenic F1 mothers. B) The distribution of CO count per BC1 progeny of each high fecundity dysgenic mother with mean and standard deviation. Asterisks denotes statistical significance by least square means (*p* < 0.05). Progeny from mother 701 had a higher average CO count than progeny from other mothers while progeny from mother 4029 exhibited a lower average CO count.

The higher recombination rate per Mb in *D. virilis* in comparison to *D. melanogaster* is due to a higher number of COs on each chromosome. In *D. melanogaster*, chromosome arms typically have zero, one, or two COs with three COs on a single chromosome arm being rare (Miller *et al*. 2016). In contrast, a chromosome with three or more COs is common in *D. virilis*, in both dysgenic and non-dysgenic directions of the cross. Chromosomes with five COs were also observed (Figure 2). CO counts per chromosome were highly similar between the progeny of dysgenic and non-dysgenic F1 females (χ^2^ 4 = 0.529, *p* = 0.97). Likewise, there was also no difference between dysgenic mothers if they were high fecundity (χ^2^ 4 = 3.70, *p* = 0.45) or low fecundity (χ^2^ 4 = 3.45, *p* = 0.49). Since per chromosome CO counts provide the data for tetrad analysis, these results indicate that dysgenesis does not modulate tetrad exchange class frequency. Therefore, we determined whether interference was altered by dysgenesis by comparing the distribution of distances between crossovers identified in the progeny of dysgenic and non-dysgenic flies. We found no differences in the distribution of crossovers for any chromosome (Mann-Whitney U test, *p* > 0.5). Overall, we find no differences in the recombination landscape between dysgenic and non-dysgenic F1 mothers in *D. virilis* at the global level. This suggests there is little to no feedback between putative activation of the DNA damage response during dysgenesis in *D. virilis* and the modulation of meiotic recombination.

**Figure 2:**
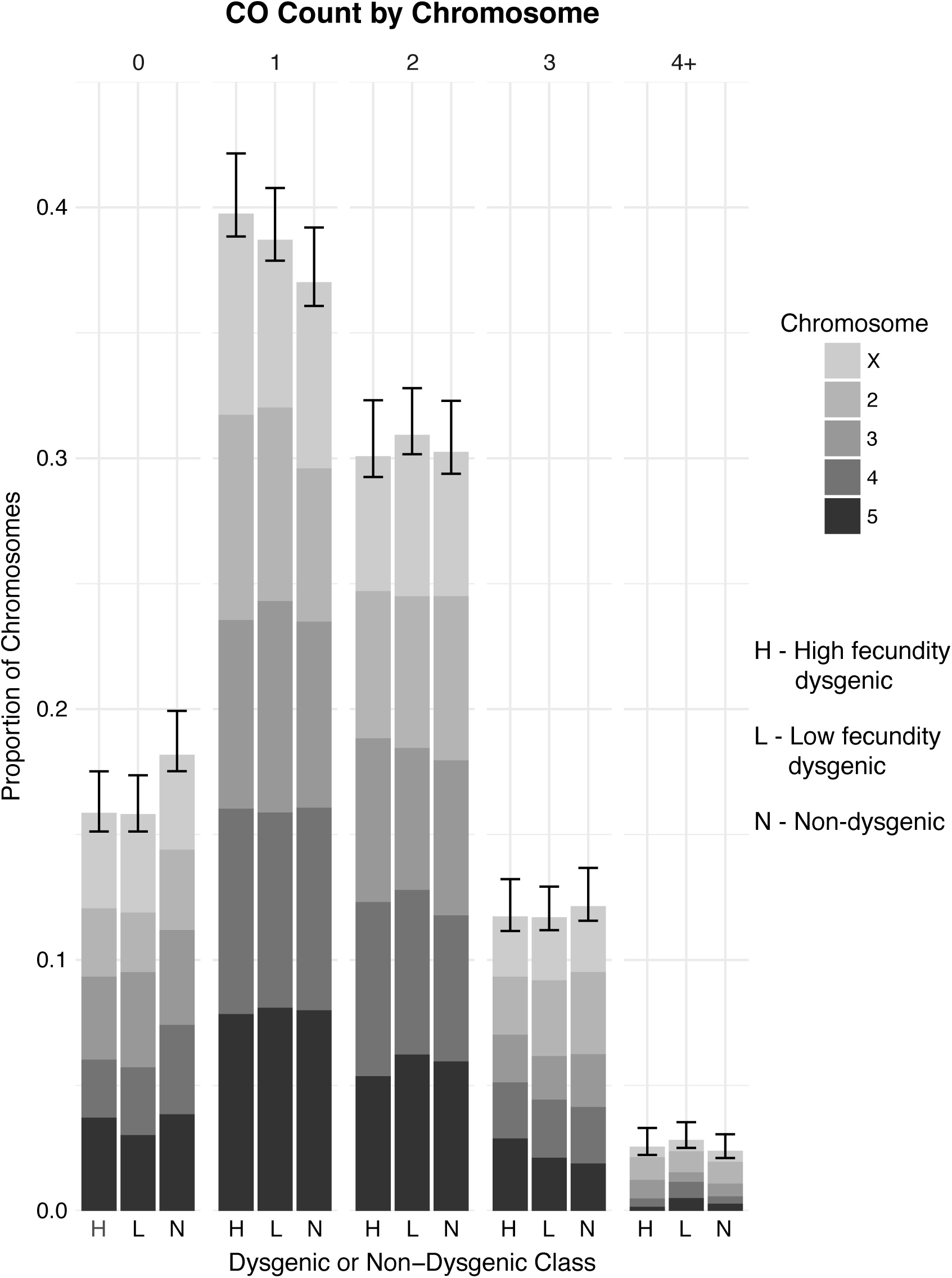
The proportion of chromosomes grouped by CO count in BC1 progeny of high fecund dysgenic, low fecund dysgenic, and non-dysgenic F1 mothers. 95% confidence intervals were calculated by sampling BC1 progeny by bootstrapping 1000 times.

We also examined the distribution of recombination events along the length of each chromosome between non-dysgenic flies, high fecundity dysgenic flies, and low fecundity dysgenic flies. There were no major changes in the distribution of recombination along the length of the chromosomes (Figure 3). The chromosomal recombination rates between all three groups are strongly correlated (Table S8). Interference plays a role in determining CO positioning.

**Figure 3:**
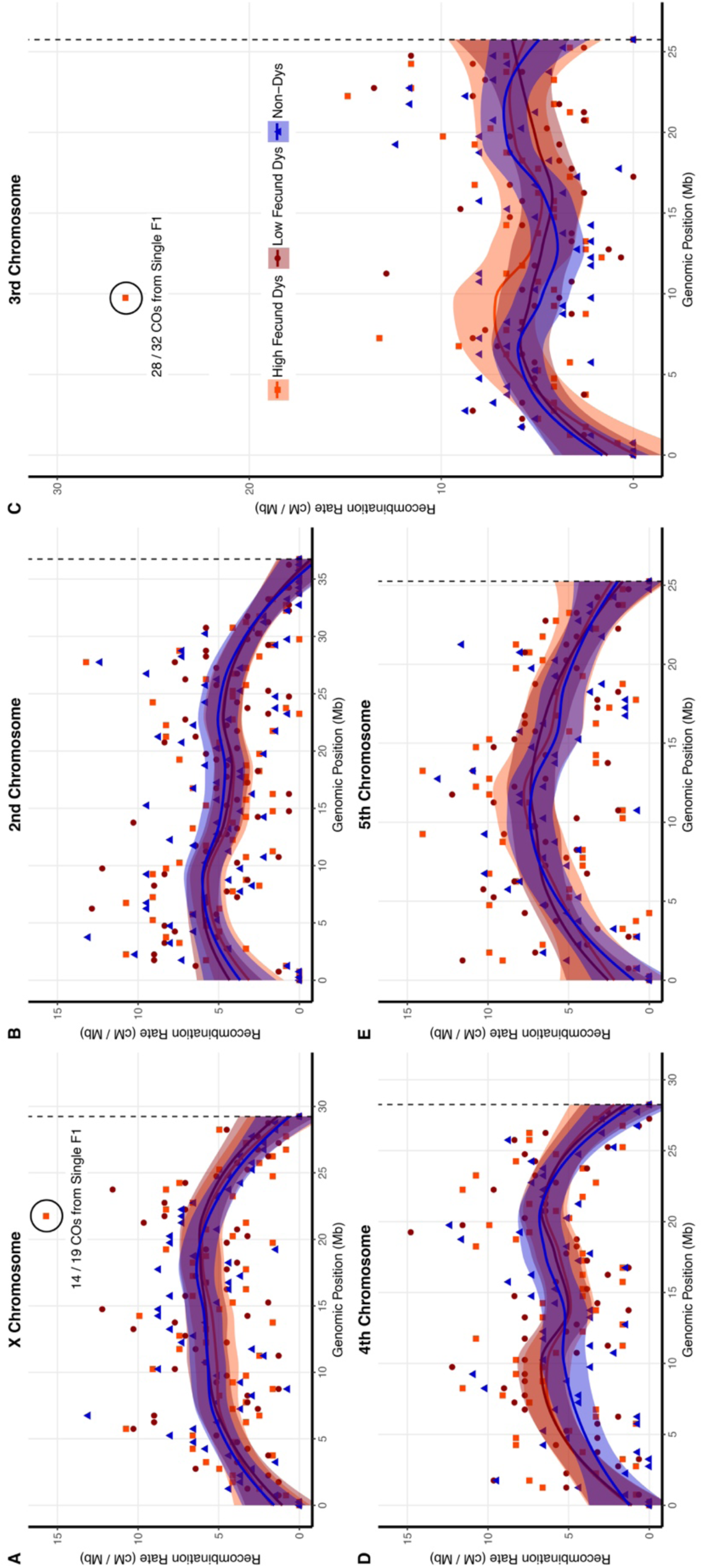
Loess smoothed splines of the recombination rate along the length of each chromosome in *D. virilis* from the telomere (left) to the centromere (right) with standard error. The dotted line represents the centromere effect of recombination suppression as recombination = 0 from the line to the end of the sequence. The rate of recombination was calculated in 500 kb intervals in F2 progeny of low fecund dysgenic, high fecund and non-dysgenic F1 mothers for the A) X chromosome, B) 2nd chromosome, C) 3rd chromosome, D) 4th chromosome, and E) 5th chromosome.

### A signature of early DNA damage and mitotic crossing over in dysgenesis

Despite observing no significant effect of dysgenesis on meiotic recombination rates, we observed several genomic regions that exhibited a much higher apparent number of COs during hybrid dysgenesis. For example, within a 500 kb region on the third chromosome, the apparent recombination rate was 26 cM/Mb, nearly twice as high as any other interval within the genome (Figure 3C, within the circled region). 32 COs were identified as arising from dysgenic F1 females compared to a single CO identified from non-dysgenic mothers. The COs in this interval provided evidence for a mitotic recombination cluster because the majority (28/32) were identified in the progeny of a single highly-fecund dysgenic F1 mother designated 5011. The mitotic recombination event identified is visible and identifiable in the genotypes of the BC1 progeny of F1 mother 5011 in comparison to the BC1 progeny of another high fecundity dysgenic female (4029) that lacks a cluster of recombination in this region (Figure 4A-B). Reciprocal CO products were observed with equal frequency among the BC1 progeny of F1 mother 5011 (χ^2^ 1 = 0.13, *p* = 0.727, Figure 4B) indicating no transmission bias among recombinant gametes. Additional unique COs were detected along the entire length of the third chromosome upstream and downstream of the recombination cluster. The high frequency of recombinants at the same location identified among this cohort of BC1 progeny likely indicates an event in the early germline of the F1 female 5011.

**Figure 4:**
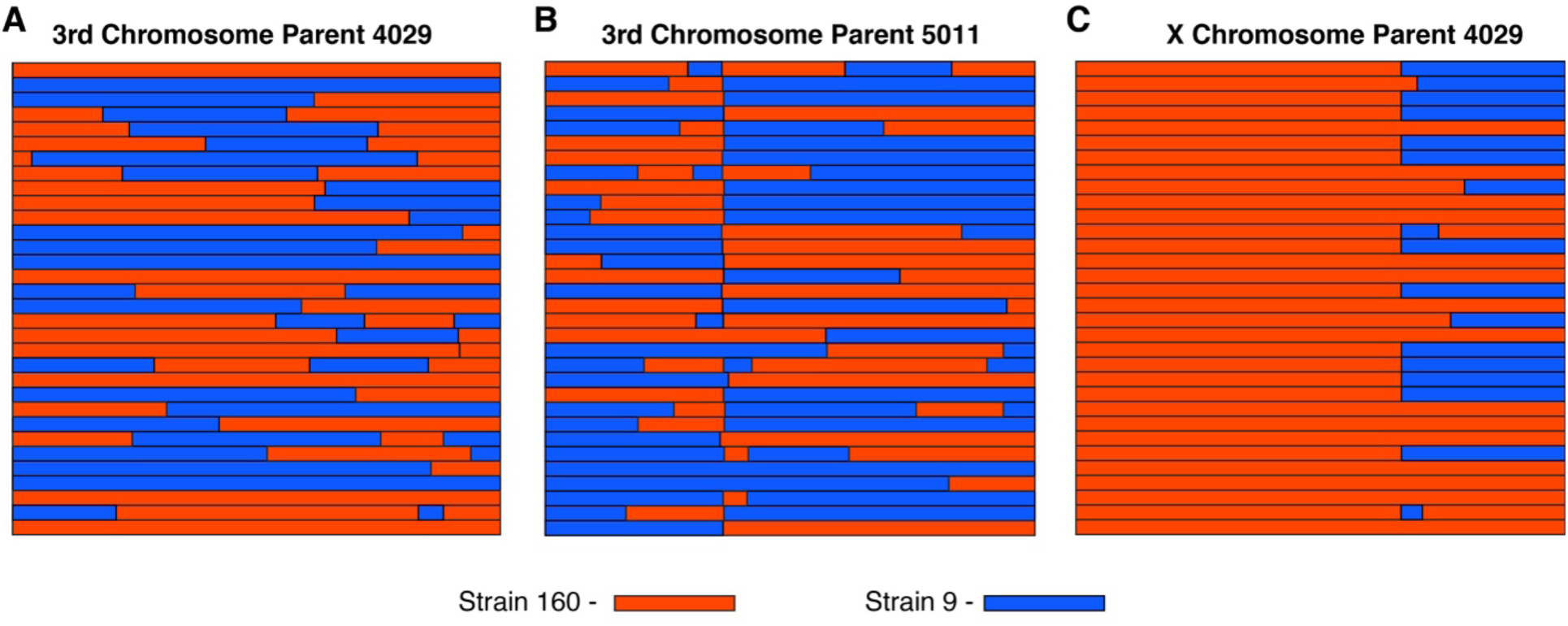
Haplotypes of BC1 progeny from a single high-fecund dysgenic mother. A) Haplotypes of the third chromosome in progeny of the 4029 F1 mother is typical of most chromosomes with no cluster of recombination. B) Haplotypes of the third chromosome in progeny of the 5011 F1 mother identify a common recombination breakpoint in most of the progeny and reciprocal products of recombination in equal frequency (Binomial test, *p* > 0.05). C) Haplotypes of the X chromosome in progeny of the 4029 F1 mother indicate a common recombination breakpoint in half of the progeny and extreme segregation distortion of the distal portion of the chromosome (227 markers 0.5 −21.4 MB, Binomial test, *p* < 1.6E-08). The proximal region of the chromosome shows no segregation distortion (86 markers 21.5 - 29.0 Mb Binomial test, *p* > 0.5).

Another recombination cluster was identified on the X chromosome, approximately 21.7 Mb from the telomere. In this region, there was an effective recombination rate of 15.7 cM / Mb (Figure 3A, within the circled region). Once again, the vast majority of COs within this 500 kb interval are part of another cluster of recombination attributed to a single highly-fecund dysgenic F1 female designated 4029 (Figure 4C). The cluster of recombination is revealed in only half of the progeny of the F1 mother 4029. Interestingly, no additional COs were detected in the portion of the X chromosome distal from the recombination event and all markers in the distal portion were heterozygous. The extreme excess of heterozygosity on the X chromosome in the BC1 progeny indicates transmission distortion of the strain 160 genotype distal from the CO from the 4029 mother (χ^2^ 1 = 32, *p* = 0.141E-08, Figure 4C). Markers proximal to the cluster of recombination show no transmission distortion (χ^2^ 1 = 0.13, *p* = 0.727, Figure 4C). Moreover, crossing over was found to occur in the proximal portion of the X. Thus, it appears that there was germline loss of heterozygosity (LOH) for the distal region of the chromosome in the germline of mother 4029.

These two clusters of recombination were identified based on their observed higher rates of recombination in the dysgenic germline. We infer the clusters are mitotic recombination hotspots because the chromosomes with the focal recombination event were exclusively found to be derived from a single F1 mother. Additional mitotic COs may be indistinguishable from meiotic recombination since such events may be rare and are only evident if the events occur early enough in development and are associated with depletion of other non-CO germline stem cells. To uncover additional evidence for other mitotic COs in our data, we screened for clusters of recombination by identifying CO events within the same 100 kb interval in four or more progeny of a single F1 mother. Using these criteria, we identified five additional candidate mitotic recombination clusters in progeny from dysgenic mothers and one additional candidate in progeny from a non-dysgenic mother (Table 4). Four of these six additional putative clusters of recombination were also associated with segregation distortion of a single genotype in a significant portion of the chromosome and no additional COs detected in the distorted region (Table 4, Figure S3).

To address whether activation of TEs during dysgenesis played a role in causing clustered mitotic recombination events, we generated a *de novo* PacBio assembly for strain 160 and determined whether regions of the 160 inducer chromosomes where recombination clusters were identified contained intact copies of TEs implicated in hybrid dysgenesis (*Penelope*, *Polyphemus*, *Helena*, *Skippy*, and *Paris*). Active versions of these TEs are absent in strain 9 and their activation during hybrid dysgenesis may induce DNA damage on the 160 chromosome for subsequent repair *via* mitotic recombination. Of these, *Paris* and *Polyphemus* are the most likely associated with chromosome breaks since they are DNA transposons capable of excision. By examining the PacBio assembly of strain 160, we found that five clusters of recombination contained an intact insertion for a TE known to be absent in strain 9 and present in strain 160 within the defined boundaries of the CO (Table 4). Three clusters were associated with *Polyphemus* elements in strain 160. One cluster was associated with a *Paris* element and a fifth cluster on chromosome X contained both elements (Table 4). To determine whether mitotic recombination events are directly associated with excision during dysgenesis, we performed PCR and sequencing on original DNA samples used for Illumina genotyping of the BC1 progeny of F1 mother 5011 with primers that flanked the *Polyphemus* insertion on chromosome 3.

Examination of the one individual that retained the strain 160 haplotype across the CO breakpoint indicated that even though there was no recombination event for that sample, excision of *Polyphemus* was identified in the lesion left by the tandem site duplication. In four recombinant samples, we confirmed that *Polyphemus* was absent. Recombination events initiated from an excision are expected to be repaired off the non-insertion chromosome and, thus we were not able to find direct evidence of an excision scar in recombinants lacking the *Polyphemus* element. Nonetheless, these results support the conclusion that this particular *Polyphemus* insertion was activated in female 5011 and this was associated with a cluster of mitotic recombination. Overall, our results suggest clusters of recombination occur more frequently in dysgenic relative to non-dysgenic females and often occur in regions containing intact copies of two DNA transposons (*Polyphemus* and *Paris*) associated with hybrid dysgenesis.

Since we identified multiple mitotic clusters of recombination in dysgenic crosses, we sought to more formally determine whether there was evidence for a statistically significant higher rate of recombination cluster formation in dysgenic females. Since the criteria for identifying a cluster was based on observing four or more individuals with a CO within a given span, it was necessary to account for variation in cohort size. We achieved this by developing a likelihood model where the probability of observing a set of chromosomes providing evidence of a cluster within a cohort was a function of the probability that a mitotic event occurs on that chromosome within an F1 female (α) and the probability of observing that event on a given chromosome (β) among the sequenced progeny. We considered two, three and four parameter models where either α or β would be the same under dysgenesis or non-dysgenesis, or there would be a unique value depending on condition. Using nested likelihood ratio tests, we find support for a three-parameter model with separate β estimates for dysgenic and non-dysgenic mothers and a shared α estimate over the two-parameter model (α = 0.12, β_Dys_ = 0.78, β_NonDys_ = 0.11, LRT, χ^2^ 1 = 51.6, *p* = 6.92E-13, Table S9). Comparing a three-parameter model where α is estimated separately to the two-parameter model is not supported (LRT, χ^2^ 1 = 3.39, *p* = 0.066, Table S9). Finally, we do not find support for a four-parameter model over the three-parameter model with separate β estimates (LRT, χ^2^ 1 = 1.88, *p* = 0.170, Table S9). Overall, these results support a similar baseline rate of occurrence of mitotic events in the dysgenic and non-dysgenic germlines.

However, the frequency of chromosomes that are transmitted with mitotic damage is greater in the dysgenic germline. Overall, we find evidence that that the total transmission rate of mitotic damage (α * β) is more than six times greater in the dysgenic germline (0.096 mitotic COs per gamete in dysgenic, 0.014 mitotic COs per gamete in non-dysgenic).

## DISCUSSION

### A high-resolution genetic map of *D. virilis*

In this study, we obtained the first high-resolution genetic map of *D. virilis*. Combined with recombination studies in *D. simulans*, *D. mauritiana*, *D. yakuba*, *D. persimilis*, *D. miranda*, *D. serrata*, *D. mojavensis*, and others (Evgen’ev 1976; True *et al*. 1996; Takano-Shimizu 2001; Staten *et al*. 2004; Kulathinal *et al*. 2008; Stevison and Noor 2010; Stocker *et al*. 2012; Smukowski Heil *et al*. 2015; Howie *et al*. 2019), our genetic map of *D. virilis* will aid future studies of the evolution of recombination in *Drosophila*. Of note is the high rate of recombination in *D. virilis* in comparison to other species, especially *D. melanogaster*. Recombination rates in species of *Drosophila* frequently peak in the middle of the chromosome arm and decrease towards the centromere and telomere (True *et al*. 1996). However, the distribution of recombination on the second, third, and fourth chromosomes in *D. virilis* resembles a bimodal distribution (Figure 3). The bimodal distribution may be the result of the exceptionally high recombination rates in *D. virilis*. When two or more crossovers on a single chromosome is common, interference preventing CO formation in close proximity would spread COs more evenly along the length of the chromosome.

### Meiotic recombination in light of hybrid dysgenesis in *D. virilis*

This study is one of the few to examine the effects of TE activity on the meiotic recombination landscape, and the first to do so using the hybrid dysgenesis syndrome in *D. virilis*. Results from previous studies of hybrid dysgenesis in *D. melanogaster* were conflicting as some found no effect on female recombination (Hiraizumi 1971; Chaboissier *et al*. 1995) while others found increases in the recombination rate (Kidwell 1977; Broadhead *et al*. 1977; Sved *et al*. 1991) or changes in the distribution of recombination (Slatko 1978; Hiraizumi 1981). In addition to reporting findings using the dysgenic syndrome of a different species, ours is also the first study to investigate the effects of hybrid dysgenesis on recombination using high-throughput genotyping rather than phenotypic markers. This allows a greater insight into the fine-scale changes in recombination rates and distribution that may previously have escaped unnoticed.

Modulation of recombination by hybrid dysgenesis may occur through different mechanisms. First, mitotic recombination would be directly initiated by double-stranded breaks (DSBs) that arise from either TE insertion or excision. Second, DSBs caused by TE activity could modulate global recombination rates through DNA damage signaling. In support of the first hypothesis, we identified clusters of recombination originating from mitotic CO events in germline stem cells (GSCs) (Figure 5). During hybrid dysgenesis, the damaging effects of transposons are often observed in the germline during early development (Engels and Preston 1979; Niki and Chigusa 1986; Sokolova *et al*. 2010). DSBs produced as an outcome of transposition may be repaired by one of several mechanisms including homologous recombination *via* mitotic crossing over. In the case of the cluster of COs on the third chromosome in F1 mother 5011, excision of a *Polyphemus* DNA transposon may have produced a DSB repaired *via* homologous recombination in the mitotic germline, in G1 prior to DNA replication, within a developing GSC (Figure 5A). In this scenario, reciprocal products of the CO would appear in all daughter cells descended from this GSC and reciprocal products would be observed, on average, in equal frequency among gametes. Alternatively, a mitotic CO may have occurred after DNA replication in G2 prior to mitosis in the germline of the 5011 mother (Figure 5B). During mitosis, the chromatids segregated according to Z segregation (Germani *et al*. 2018; Pimpinelli and Ripoll 1986) such that reciprocal CO products were transmitted to one daughter cell while the other daughter cell would have received the non-CO chromatids. Other GSCs must have been retained within the 5011 mother because there are several progeny without the common CO product. However, the limited number of progeny lacking either of the reciprocal CO chromatids indicates a depletion of other intact GSCs. We attribute this to an early crisis in GSC survival due to hybrid dysgenesis, followed by re-population of the GSCs from descendants of a single cell carrying the CO chromatids. GSCs marked with the mitotic CO were thus able to recover and rescue fertility after hybrid dysgenesis in the high fecundity female. This is consistent with the observation that hybrid dysgenesis is associated with an early phase of germline depletion (Sokolova *et al*. 2013).

**Figure 5:**
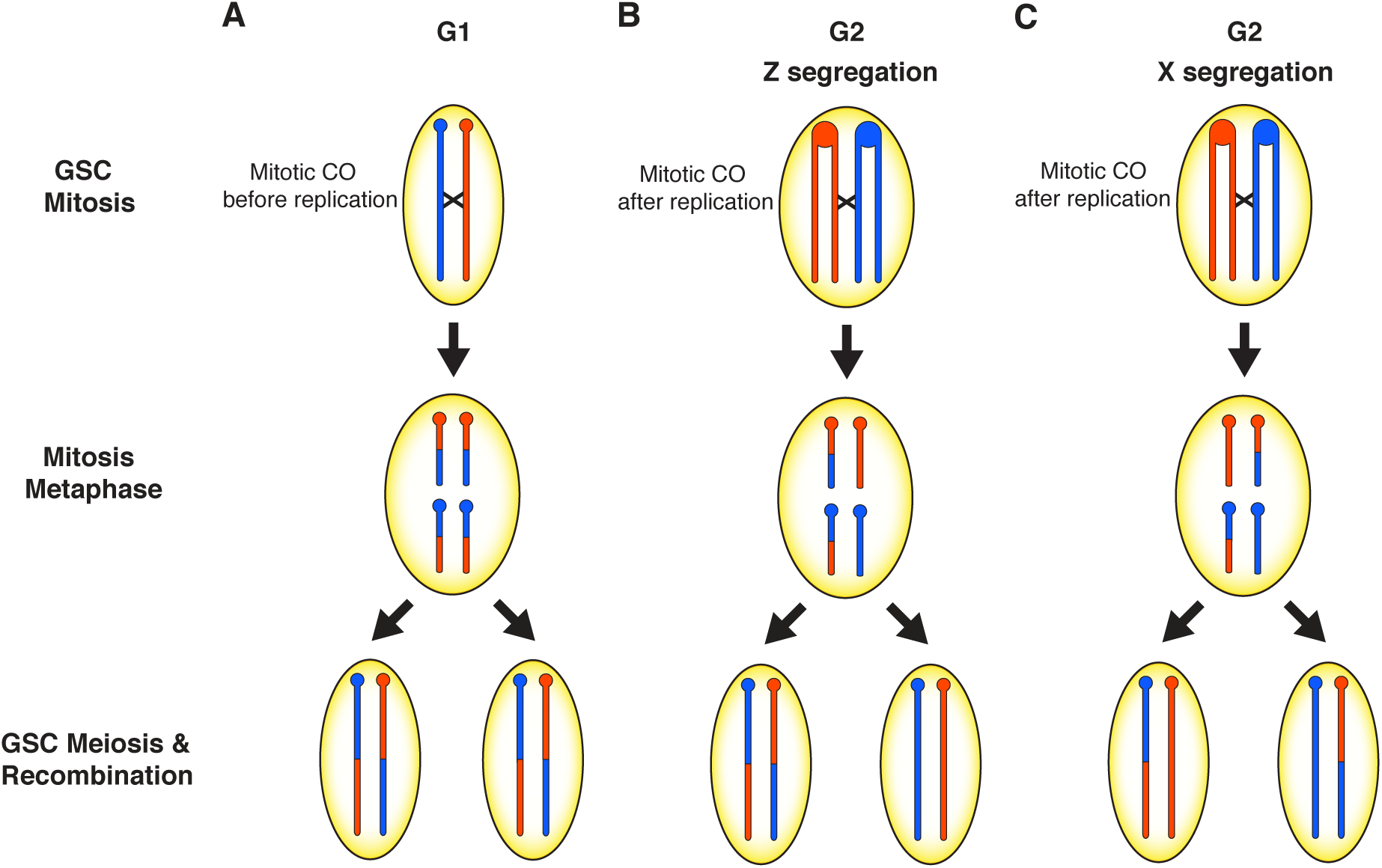
Models to explain the clusters of recombination on the third and X chromosomes in the progeny of two high fecundity dysgenic mothers. In the 5011 F1 female, a mitotic crossover (CO) either occurred A) prior to DNA replication in the early developing germline resulting in two daughter cells with the CO or B) after DNA replication, followed by a pattern of Z segregation so that one daughter cell has both CO chromatids. Oocytes produced by these germline stem cells will transmit the CO as reciprocal products as seen if the CO occurred in G1. C) A mitotic CO in the 4029 F1 female occurred after DNA replication in the developing germline and each daughter cell received one CO chromatid and one non-CO chromatid according to a pattern of X segregation. This results in a loss of heterozygosity (LOH) in the distal portion of the chromosome to resemble segregation distortion and recombination events are not detectable.

A second mitotic CO event on the X, with different consequences, likely occurred in the early developing germline of the 4029 mother (Figure 5C). In this case, there was apparent loss-of-heterozygosity (LOH) distal from the CO. This mitotic CO event would have occurred after DNA replication in G2, with a pattern of X segregation (Germani *et al*. 2018; Pimpinelli and Ripoll 1986) resulting in each daughter cell receiving one chromatid with the CO and one without and LOH distal from the CO. This LOH would be responsible for failure to detect additional meiotic COs derived from the homozygous distal region. The complete transmission distortion of the distal region suggests severe depletion of GSCs with the reciprocal mitotic CO products. Again, this is consistent with a severe reduction of GSCs, followed by re-population from even a single GSC and restoration of fertility in the high fecundity female. Interestingly, the bounds of the mitotic CO derived from the 4029 mother do not contain intact dysgenesis-associated TEs nor any other intact TEs in the strain 160 genome. This mitotic CO may therefore have been the product of a new TE insertion in the genome of the 4029 mother. LOH is also observed among several clusters of recombination and most of these clusters are associated with either *Polyphemus* or *Paris* DNA transposons (Table 4, Figure S3). LOH *via* mitotic recombination is observed after DNA damage or chromosomal breakage in cancer cells (Lasko 1991) and in yeast recombination studies (Symington *et al*. 2014). The greater number of mitotic recombination events with segregation distortion observed in our data is consistent with previous observations of non-random segregation of chromatids in clonal analysis; chromatids involved in mitotic exchange are more likely to segregate into separate daughter cells (X segregation) than the same daughter cell (Z segregation) in mosaic analyses (Germani *et al*. 2018; Pimpinelli and Ripoll 1986) Likewise, segregation distortion is frequently observed during hybrid dysgenesis (Kidwell and Kidwell 1975; Kidwell *et al*. 1977). Our study therefore links segregation distortion *via* mitotic recombination and subsequent LOH within female germlines as a result of hybrid dysgenesis.

The observed number of mitotic CO events identified in dysgenic progeny is interesting because the crossing over pathway is least likely to repair non-programmed DSBs (Preston *et al*. 2006). Mitotic COs only account for ∼1% of all COs detected in our dataset and contribute minimally to the genetic map length (Figure S4). Interestingly, the mitotic exchange rate is similar to the rates of male recombination under *P* element hybrid dysgenesis (Kidwell *et al*. 1977; Preston *et al*. 1996). Other pathways of DNA damage repair including non-homologous end joining and single-strand annealing are probably more common but undetectable in our assay. Rates of mitotic crossing over may also be higher than what we estimated since many mitotic COs would not meet our criteria for classifying candidate clusters because many dysgenic mothers produced fewer than four offspring.

Overall, despite evidence for the mitotic recombination associated with DNA transposon excision, we found no major differences in the distribution and frequency of meiotic recombination in *D. virilis* under hybrid dysgenesis, suggesting that DNA damage signaling during dysgenesis does not modulate meiotic recombination. The DNA damage response has a critical role in regulating meiotic recombination (Sjöblom and Lähdetie 1996; Lu *et al*. 2010; Joyce *et al*. 2011). DNA damage response regulators such as *p53* and *chk2* also influence the fate of GSCs during hybrid dysgenesis (Tasnim and Kelleher 2018). The incomplete penetrance of hybrid dysgenesis in *D. virilis* may arise from cell to cell variation in the total amount of DNA damage or in stochastic variation in the DNA damage response. However, we found no differences in recombination rates between dysgenic flies with minimal germline atrophy (high fecundity) and severe germline atrophy (low fecundity). This suggests that DNA damage signaling activated by dysgenesis does not modulate meiotic recombination. This is likely due to the fact that most TE activity happens in GSCs during early development (Engels and Preston 1979; Sokolova *et al*. 2010). By the onset of meiosis, the harmful effects of TE activity during dysgenesis may have likely subsided. In *D. virilis*, TE suppression is restored by adulthood in dysgenic progeny *via* re-establishment of asymmetric piRNAs and the negative impacts of dysgenesis disappear in subsequent generations (Erwin *et al*. 2015). This suggests that TEs likely produce few DSBs during meiosis in the *D. virilis* hybrid dysgenesis model. We thus conclude that the timing of transposition is an important factor that determines how TEs influence recombination. In the future, it will be worth investigating whether recombination is also robust to transposition that occurs closer to the initiation of meiotic recombination.

## Acknowledgements

We thank John Kelly, Stuart Macdonald, Michael Evgen’ev, Sergei Funikov, members of the Robert Unckless and Amanda Larracuente labs, and anonymous reviewers for helpful comments and suggestions. Peter Andolfatto and Audrey Lamb provided guidance on the multiplex shotgun genotyping protocol and Annemarie Chilton produced the custom Tn5 enzyme. Enzyme production and development of the library generation protocol was in part supported by NIH R01 OD010974 to Stuart Macdonald (KU) and Tony Long (UC Irvine). We also thank Jennifer Hackett and the University of Kansas Genome Sequencing Core (NIH COBRE P20GM103638) and Christina McHenry and the University of Michigan Advanced Genomics Core for library prep and sequencing. We also thank the KU Center for Research Computing, especially Riley Epperson, and the Georgia Advanced Computing Resource Center for computational resources. This work was supported by the University of Kansas, the University of Georgia, support from COBRE P20GM103638 and NSF Award 1413532 to JB.

